## Supplementary material for "The pharmacoepigenomic landscape of cancer cell lines reveals the epigenetic component of drug sensitivity"

### This PDF file includes:

Figure S1 to S7
Legends for Table S1 to S2

### Other supplementary materials for this manuscript include:

Table S1 to S2

**Table S1: Enrichment of drug classes in dDMRs.** Summary statistics of hypergeometric tests for drug class enrichments. The table contains drug class annotations, the associated hypergeometric test p-value testing for either enrichment or depletion, the adjusted p-value using the Benjamini-Hochberg false discovery rate correction and the tested cancer type.

**Table S2: 802 dDMRs, 377 short-listed dDMRs, 58 tgdDMRs and 19 tgdDMRs with protein-protein interaction networks.** Summary statistics and annotations of identified dDMRs and tgdDMRs, annotated by drug and its putative drug target, the cancer type, the genomic region, its functional region and exact position, the adjusted p-value from the discovery cohort, the proximal associated gene in cancer cell lines and primary tumours, the adjacent gene, the nature of the correlation between the methylation and expression in both cancer cell lines and primary tumours, the associated derived mechanism, the association p-value between the proximal gene expression and drug response, the number of sites in the respective region, information about a potential DHS, enhancer, an associated signalling network, the cancer gene information, its validation status in the CTRP and CCLE cohorts and associations of the their methylation with genetic alterations, CRISPR knockout screens and LINCS drug signatures.

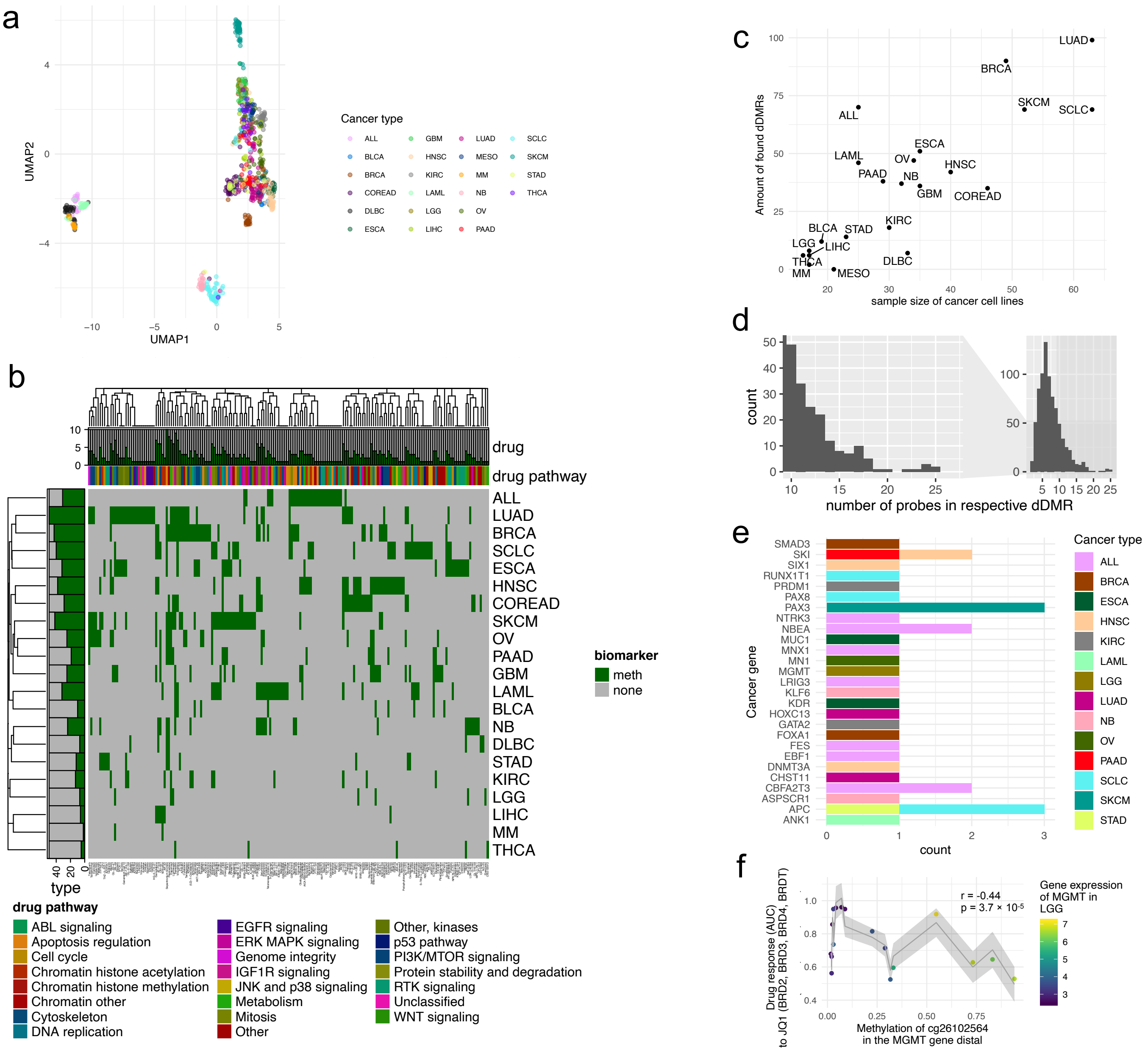


**Figure S1: Additional characterisations of dDMRs.** (**a**) The UMAP dimensionality reduction for gene expression patterns of cell lines in GDSC. (**b**) A heatmap of drugs that showed at least one dDMR across the screened compounds and cancer types. (**c**) Scatter plot showing sample size of each cancer type against found dDMRs across all screened drugs. (**d**) A histogram of the number of sites in each dDMR. (**e**) Set of 27 dDMRs proximal to cancer genes. (**f**) dDMR in *MGMT* for response to JQ1 in low-grade glioma (LGG).

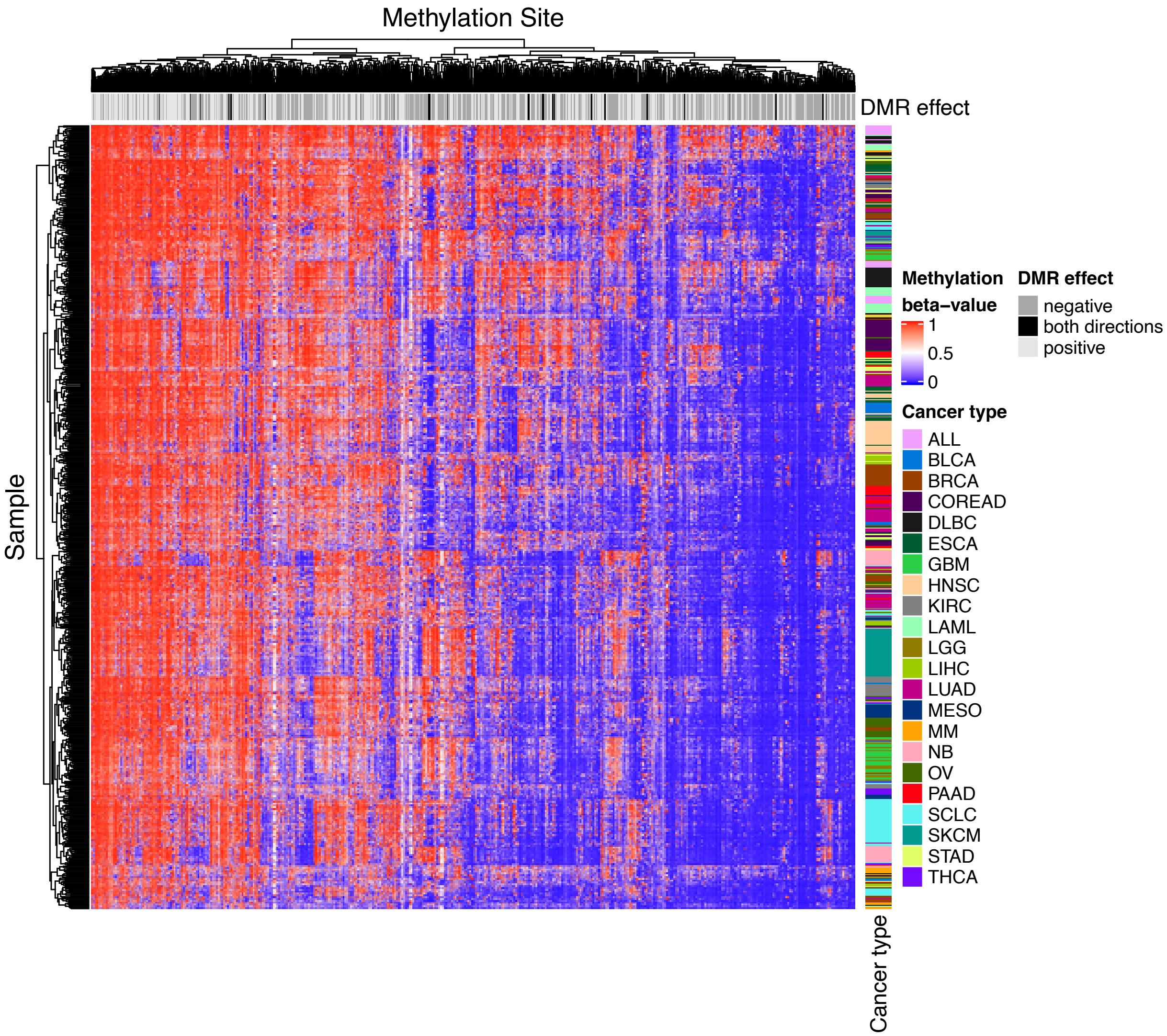


**Figure S2: Methylation pattern of dDMRs.** Heatmap of dDMR methylation across cancer types.

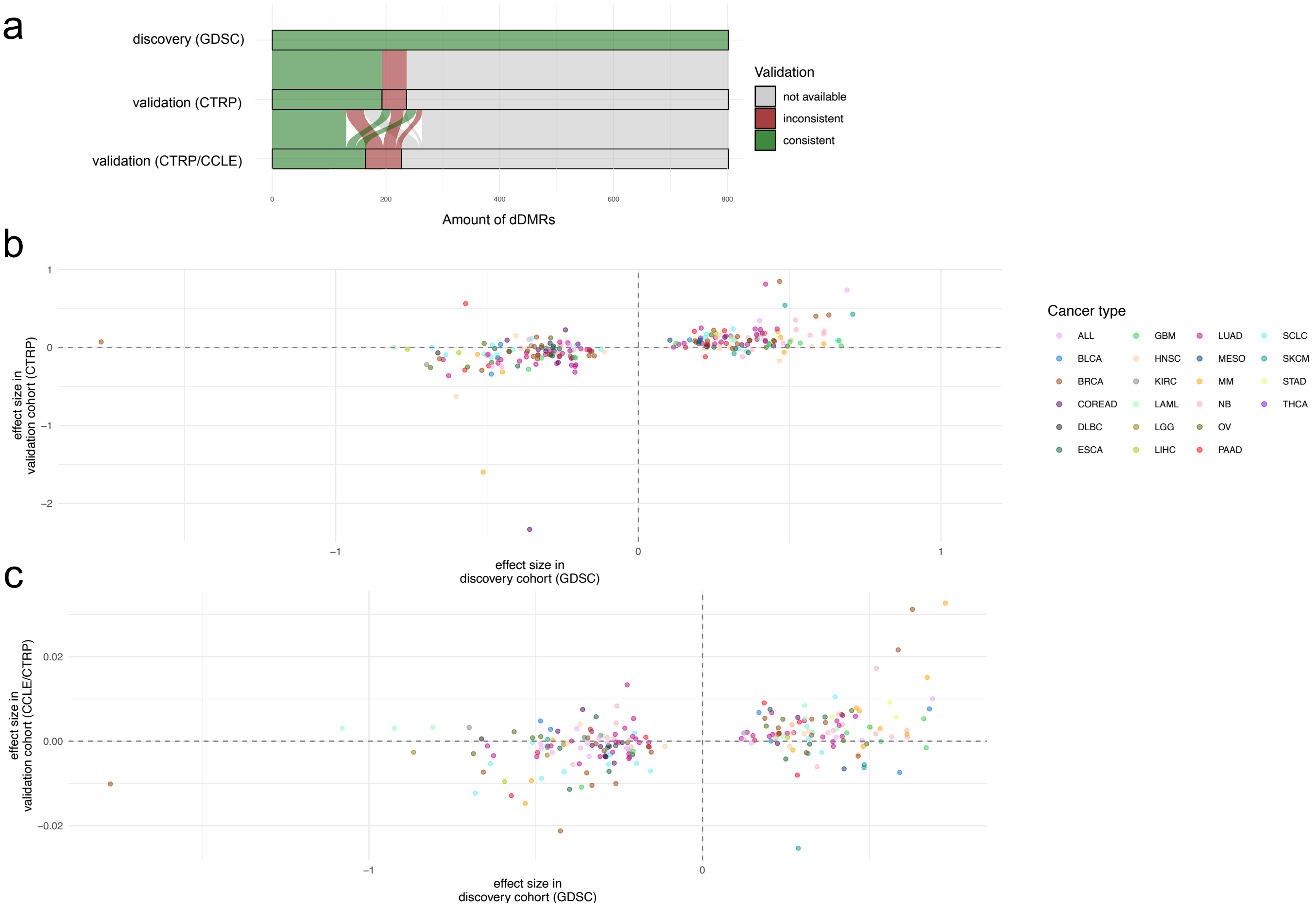


**Figure S3: Scatter plot of effect sizes from validation cohorts.** (**a**) Consistency of effect sizes of 802 dDMRs from the GDSC discovery cohort validated in either the CTRP and CCLE datasets. (**b**) Effect sizes of dDMRs validated with the CTRP drug response HTS. (**c**) Effect sizes of overlapping dDMRs with the CCLE RRBS and CTRP data.

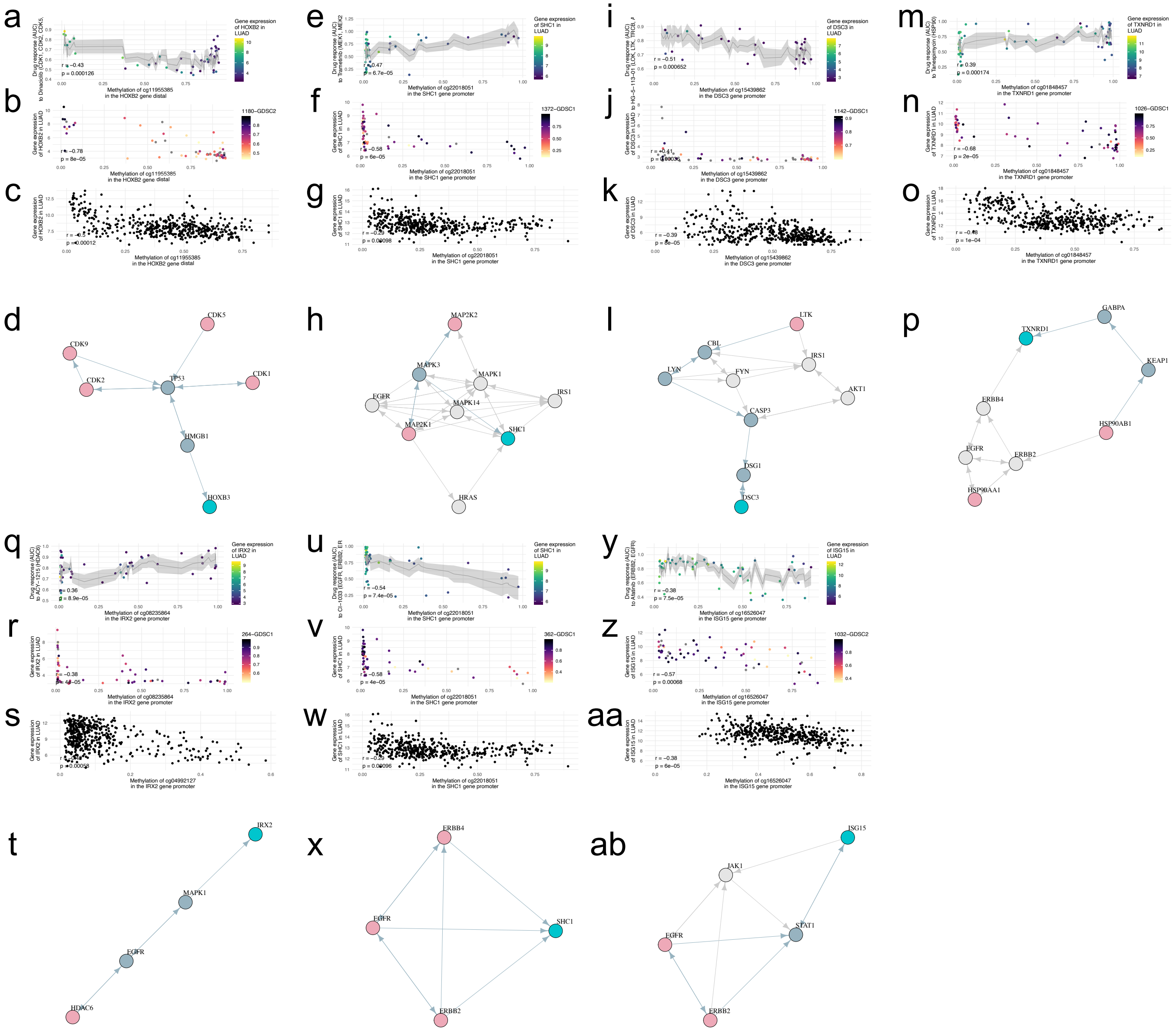


**Figure S4: tgdDMRs in LUAD.** (**a**)-(**ab**) Correlation between drug response quantified by area-under-the-curve, DNA methylation and gene expression plus the corresponding protein-protein interaction network between putative drug target (pink) and tgdDMR-associated gene encoding protein (light-blue). In the graph, nodes that are traversed with a shortest path are highlighted by the blue-grey colour among the alternative paths.

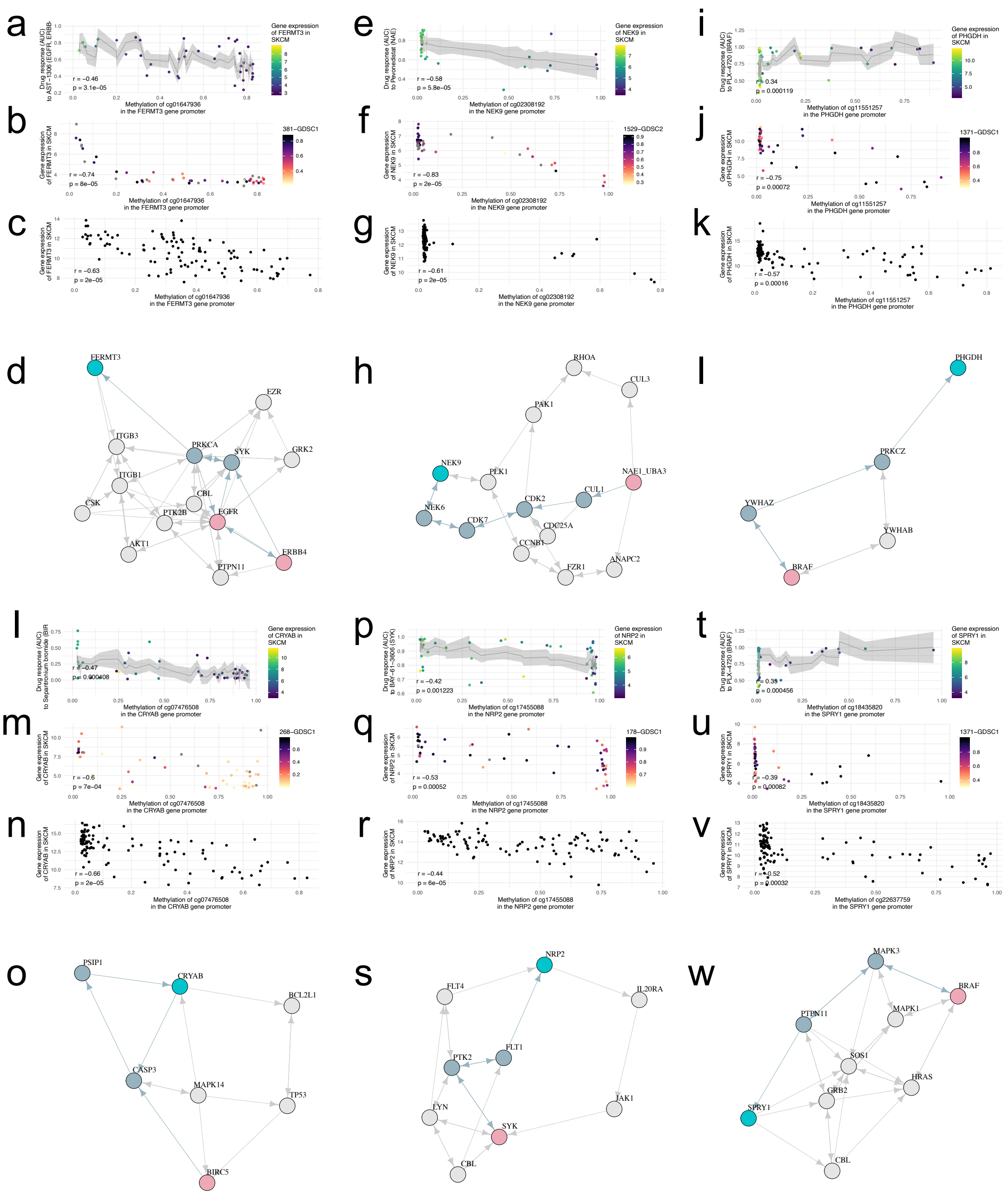


**Figure S5: tgdDMRs in SKCM.** (**a**)-(**w**) Correlation between drug response quantified by area-under-the-curve, DNA methylation and gene expression plus the corresponding protein-protein interaction network between putative drug target (pink) and tgdDMR-associated gene encoding protein (light-blue). In the graph, nodes that are traversed with a shortest path are highlighted by the blue-grey colour among the alternative paths.

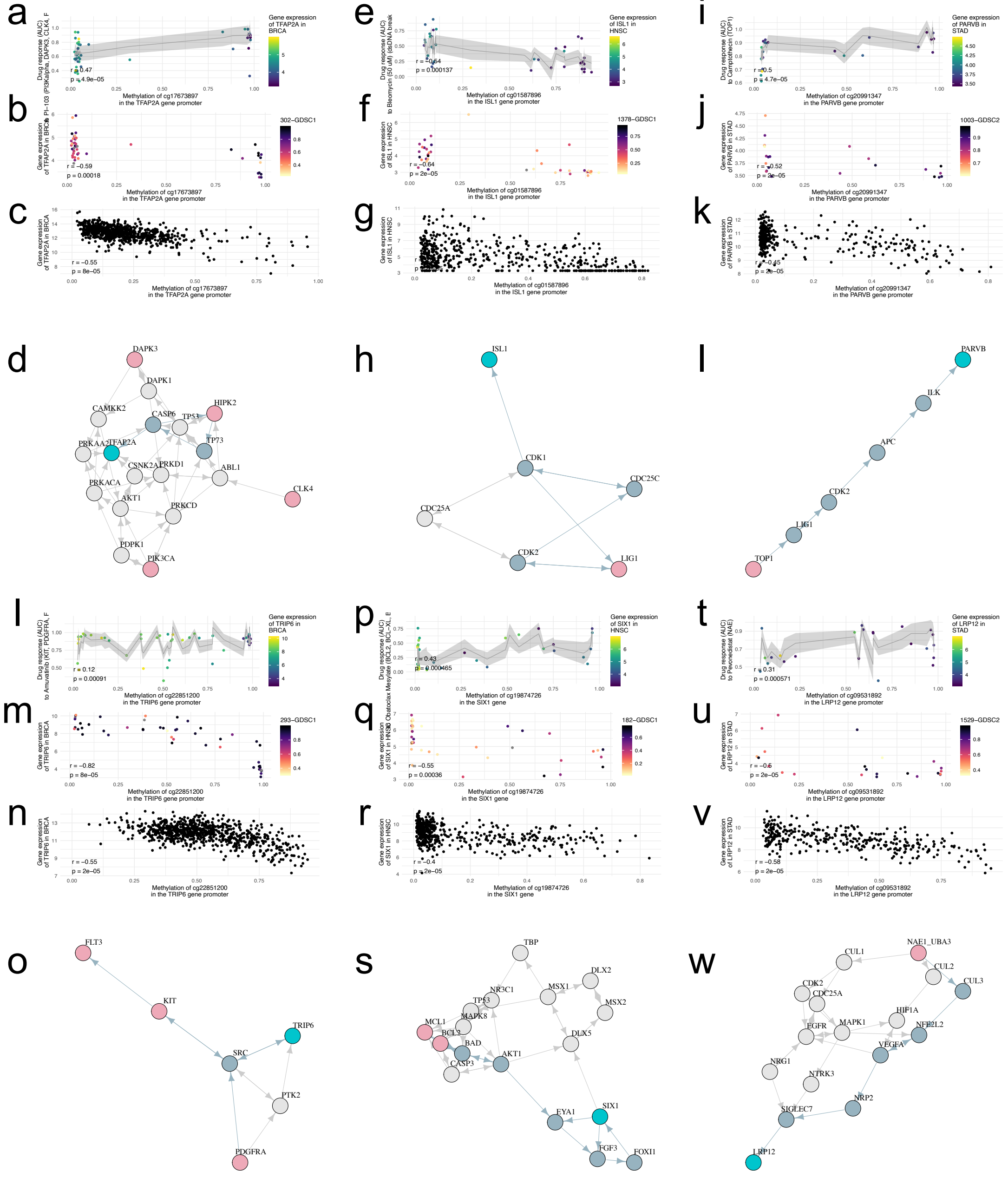


**Figure S6: tgdDMRs in BRCA, HNSC and STAD.** (**a**)-(**w**) Correlation between drug response quantified by area-under-the-curve, DNA methylation and gene expression plus the corresponding protein-protein interaction network between putative drug target (pink) and tgdDMR-associated gene encoding protein (light-blue). In the graph, nodes that are traversed with a shortest path are highlighted by the blue-grey colour among the alternative paths.

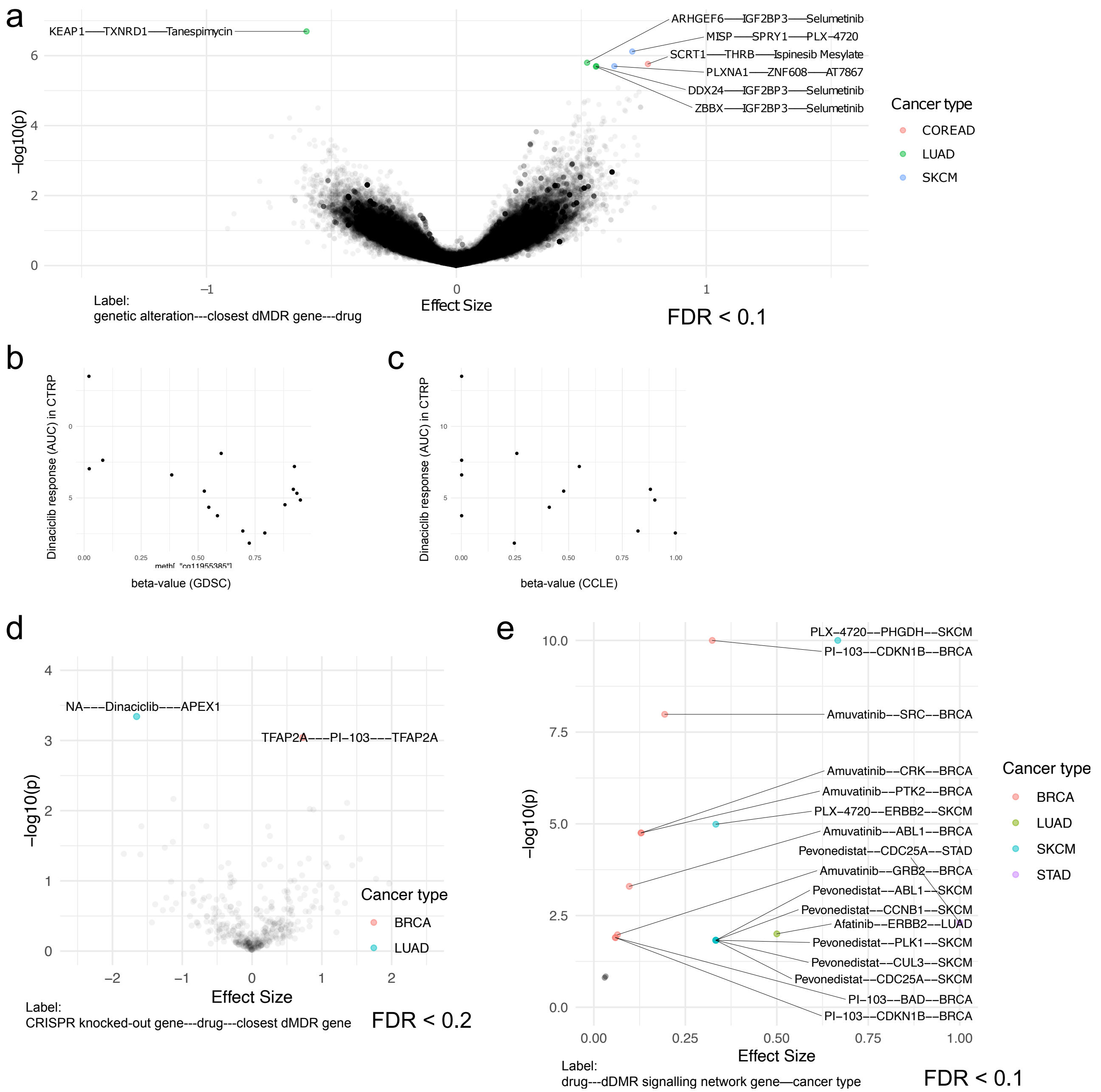


**Figure S7: tgdDMRs in the context of genetic alterations, CRISPR screens and drug signatures.** (**a**) A volcano plot summarising associations between tgdDMRs and somatic mutations in cancer cell lines. (**b**,**c**) Scatter plot for drug response to dinaciclib (AUC) in the CTRP validation set with the HumanMethylation450 BeadChip array in GDSC and reduced representation bisulfite sequencing in the CCLE independent dataset. (**d**) A volcano plot summarising associations between tgdDMRs and CRISPR knockout screens of the genes associated with tgdDMRs and their signalling network neighbourhood. (**e**) A volcano plot summarising enrichments of genes associated with tgdDMRs and their signalling network neighbourhood in the LINCS drug signatures for the matching compound and cancer type.
